# TACI: an ImageJ plugin for 3D calcium imaging analysis

**DOI:** 10.1101/2021.09.28.462182

**Authors:** Alisa A. Omelchenko, Hua Bai, Sibtain Hussain, Jordan J. Tyrrell, Lina Ni

## Abstract

Research in the field of neuroscience has evolved to use complex imaging and computational tools to extract comprehensive information from data sets. Calcium imaging is a widely used technique that requires sophisticated software to obtain reproducible results, but many laboratories struggle to adopt computational methods when updating protocols to meet modern standards. Difficulties arise due to the lack of computational knowledge and paywalls for software. In addition, most calcium imaging analysis approaches ignore motion on the z-axis. Here, we described a workflow to use ImageJ to analyze 3D calcium imaging. We applied TrackMate, an open-source ImageJ plugin, to track neurons in the lateral (x/y) direction, detect regions of interest (ROIs), and extract fluorescence intensities. To track motion on the z-axis, we developed a new ImageJ plugin, TrackMate Analysis of Calcium Imaging (TACI). For neurons appearing on multiple z-positions, maximum fluorescence values were identified to represent neurons’ intensities of corresponding z-stacks. This workflow does not require coding ability, avoids human bias, and increases reproducibility. We validated this workflow using fly larval thermosensitive neurons that displayed movements in all directions during temperature fluctuation and a 3D calcium imaging dataset acquired from the fly brain.

## Introduction

The level of intracellular calcium is a precise marker for neuronal excitability. Calcium imaging measures the changes in intracellular calcium to understand neuronal activities [1]. Studies in neuroscience have increasingly used this method due to the development of techniques for measuring intracellular calcium concentration, including genetically encoded calcium indicators (GECI), such as GCaMP [2, 3], which can be noninvasively expressed in specific sets of neurons through genetic approaches. The lower costs of lasers and microscope components have also increased the use of calcium imaging [4]. Importantly, calcium imaging allows for recording and studying single neurons as well as large neuron populations simultaneously in freely moving animals [5].

Nevertheless, the analysis of calcium imaging data is challenging because (1) it involves tracking changes in fluorescence of individual cells over time, (2) the fluorescence signal intermittently disappears or reappears with neuronal responses, and (3) neurons may shift in all directions, specifically in and out of a focal plane or appearing on multiple planes [4, 6]. Many laboratories manually perform calcium imaging analysis, which is time-consuming and becomes impractical as the length of recordings and the number of neurons increases. Manual analysis also introduces operator bias, is prone to error, and affects the replicability of experiments.

Various software has been developed to accelerate the process of analyzing calcium imaging and increase its reproducibility. Previously, software was designed in a limited experimental context making it difficult for other laboratories to adopt. Recent efforts to meet modern standards for software sharing have led to the development of several tools that can consistently analyze calcium imaging data across different groups [7–19]. However, many of these tools still require programming knowledge and/or depend on commercial software. Lack of programming knowledge and software paywalls deter researchers from adopting these methods. Moreover, most of these tools focus on correcting x/y motion and ignore motion on the z-axis [6]. Thus, there is a need for an alternative method to reproducibly analyze 3D calcium imaging that focuses on neurons appearing on multiple z-planes and exhibiting z-drift. Ideally, this tool will use open-source software that does not require programming ability so that most laboratories can readily adopt it.

TrackMate is an open-source ImageJ plugin for tracking single particles [20]. It has been widely used to track particles in various biological studies involving live-cell imaging, including calcium imaging [11, 20–22]. Generally speaking, calcium imaging analysis includes three steps: motion correction, region of interest (ROI) detection, and signal extraction [4, 12]. All three steps can be automated in TrackMate, thus significantly reducing operator bias and increasing reproducibility in calcium imaging analysis. Importantly, it is an ImageJ plugin and does not require coding ability.

Here, we developed a new ImageJ plugin, TrackMate Analysis of Calcium Imaging (TACI), to analyze 3D calcium imaging data. First, TACI organizes 3D calcium imaging data by z-positions. Then, TrackMate is applied to track x/y motion, define ROIs, and extract fluorescence intensities on each z-plane. TACI is designed to correct motion on the z-axis. Currently, z-drift is corrected by (1) using maximal or mean projection intensities [23, 24], (2) extracting fluorescence intensities from 3D ROIs [8, 11, 25], and (3) adopting maximum values across z-stacks [26]. TACI identifies the maximum value of a z-stack and uses it to represent a cell’s intensity at the corresponding time point. This workflow is suited to analyze 3D calcium imaging with motion in all directions and/or with neurons (fully) overlapping in the lateral (x/y) direction but appearing on multiple z-planes. 3D calcium imaging datasets from fly larval thermosensitive neurons and mushroom neurons in the brain were used to validate this workflow. Of note, both TACI and TrackMate are open-source ImageJ plugins and do not require any computing knowledge.

## Results

### A workflow of 3D calcium imaging analysis

In this study, we developed a new ImageJ plugin, TrackMate Analysis of Calcium Imaging (TACI), and described a workflow to use TACI to track z-drift and analyze 3D calcium imaging that pinpointed responses of individual cells appearing in multiple z-positions (Movie S1). This analysis process included three steps (Fig 1). First, TACI ORGANIZE function organized 3D calcium imaging .tif data by z-positions. Images from the same z-position were saved in one folder. TACI ORGANIZE function could grayscale these images when needed. If image names were not compatible with TACI ORGANIZE function, TACI RENAME function could convert image names to the required structure.

**Fig 1.**
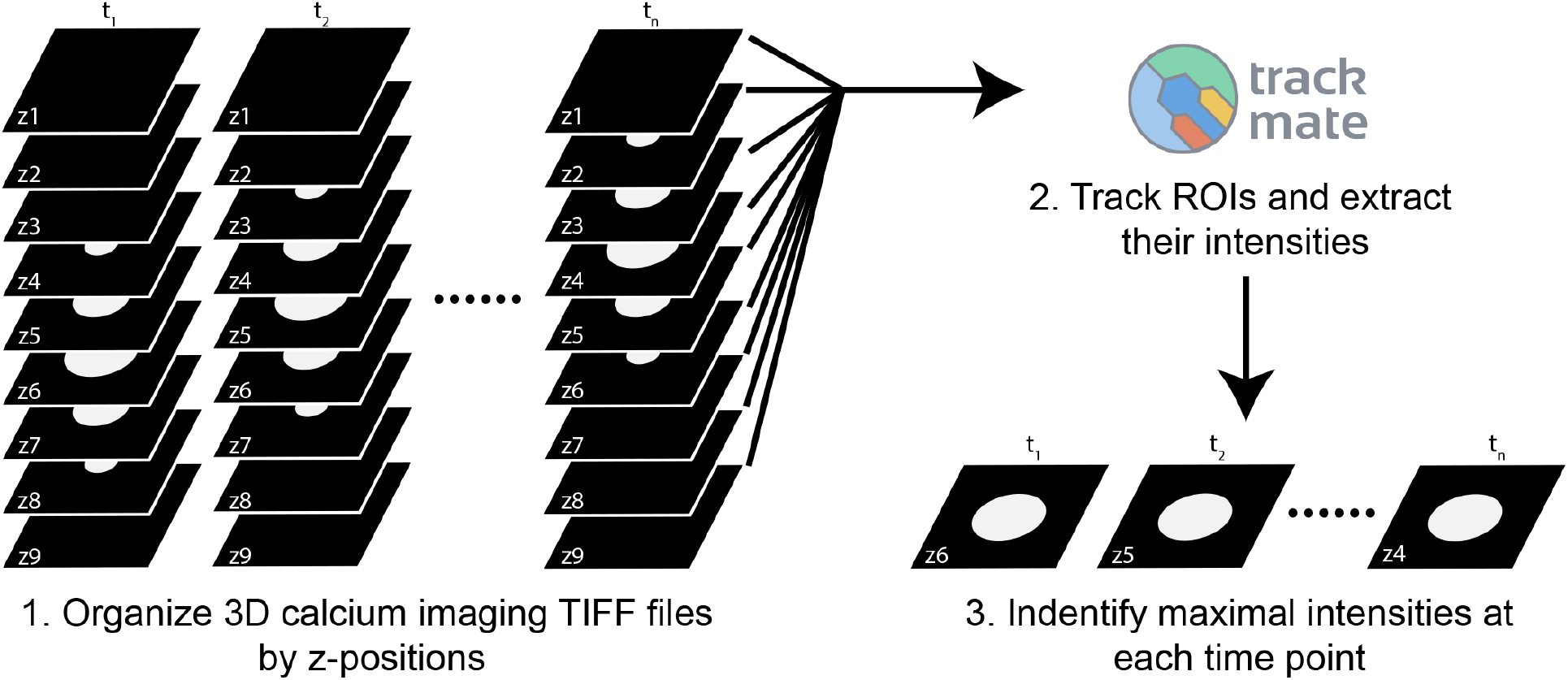
Workflow of using TACI to analyze 3D calcium imaging. This workflow includes three steps. Step 1. TACI organizes 3D calcium imaging .tif files by z-positions. Images from the same z-position are saved in the same folder. If necessary, TACI can first rename images to the required naming structure and grayscale them. Step 2. Detect and track ROIs and extract their intensities. Images from the same z-positions are analyzed as a stack. This study uses TrackMate to accomplish this step. Step 3. TACI creates a list including every time point and fills the mean intensities into the corresponding time points, identifies the maximum value of each z-stack, subtracts the background, and calculates ΔF/F_0_. TACI also calculates the average of ΔF/F_0_ of multiple cells if needed.

Second, ROIs were detected and tracked, and their fluorescence intensities were extracted on every z-plane. We used an ImageJ plugin, TrackMate, to accomplish this step. TrackMate combined the functions of ROIs’ detection and tracking and extraction of their intensities [20, 22]. Images from the same z-position were opened in order by ImageJ as one stack. Multiple TrackMate parameters were recommended to be adjusted to get optimal results, including (1) using DoG or LoG detector (Fig S1A), (2) changing the blob diameter, threshold, and median filters (Fig S1B), (3) setting filters to remove some, if not all, irrelevant signals (Fig S1C), (4) changing linking max distance, gap-closing max distance, and gap-closing max frame gap (Fig S1D), and (5) exporting all spots statistics (Fig S1E). When using the same parameter settings, fluorescence intensities extracted by TrackMate were consistent from one computer or operator to another and therefore reproducibility was increased (Table S1).

Last, for every cell of interest, TACI EXTRACT function sorted mean intensities by the corresponding time points, identified maximum values of each z-stack, subtracted the background, and calculated ΔF/F_0_. TACI MERGE function calculated the average of ΔF/F_0_ of multiple cells.

### The maximum value is a good representative of a cell’s intensity

We used simulated cells to justify that the maximum value is a good representative of a cell’s intensity. Three cells (3D spheres) of different brightness were created, and z-stacks were simulated (Fig 2A). These z-stacks had the same z-distance and z-position. A sphere’s volume and filled intensity were multiplied to create its ground truth intensity. TrackMate extracted cell intensities to determine the maximum intensity value of every cell. The ratio of the maximum value and the ground truth intensity was then calculated. We discovered that the ratio was the same for cells of various brightness. Additionally, z-stacks were built using spheres with various z-distances (Fig 2B) or z-positions (Fig 2C). The ratio of the maximum value and the ground truth intensity was kept the same. These data suggest that the maximum value is a good representative of a cell’s intensity.

**Fig 2.**
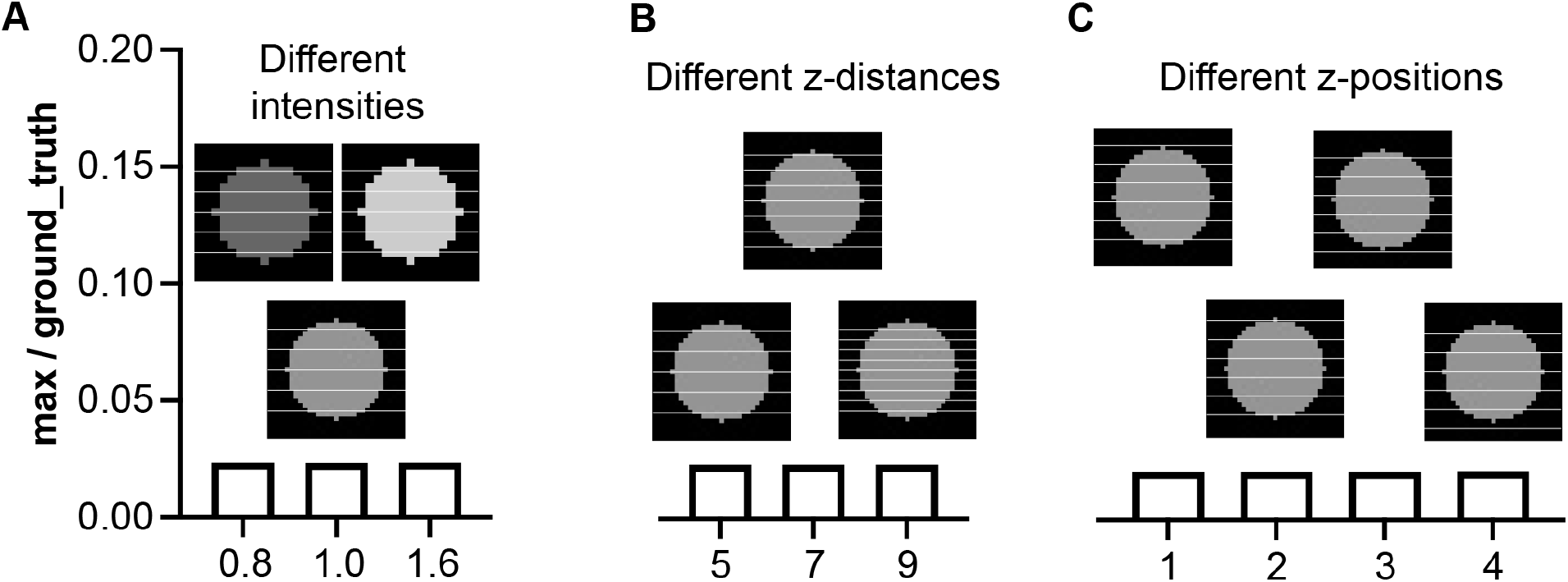
The maximum value is a good representative of a cell’s intensity. (A) z-stacks are created from simulated cells with different intensities. (B) z-stacks with different z-distances are simulated. (C) z-stacks with different z-positions are simulated. max: the maximum value of a z-stack. ground_truth: the product of the simulated cell’s volume and its filled intensity.

An alternative way to represent a cell’s intensity is to get the total of intensities from all z-positions (Fig S2). We calculated the ratio of the sum value and the ground truth intensity. We found that the ratio increased with decreasing z-distances (Fig S2B), although it remained constant if spheres had varied brightness (Fig S2A) or z-stacks were at different z-positions (Fig S2C). Of note, simulated data were obtained under ideal conditions. According to our experience, stronger signals were easier to detect and extract. Weak signals tended to be ignored, and thus it was more challenging to precisely extract their intensities. Since weak signals only affect the sum value, but not the maximum value, the maximum value indicates a cell’s intensity more properly.

### Fly larval cool neurons respond to temperature changes

We validated this method using the calcium changes to temperature fluctuations in fly larval cool neurons. A genetically encoded calcium indicator, GCaMP6m [27], was expressed in larval cool neurons by *Ir21a-Gal4* [28]. When exposed to approximately 27°C, the neurons had a low intracellular calcium levels (Fig 3A and 4A). When the temperature was decreased to approximately 10°C and held, the intracellular calcium levels rapidly increased and sustained (Fig 3B and 4A). The calcium levels rapidly dropped when the temperature was increased (Fig 4A).

**Fig 3.**
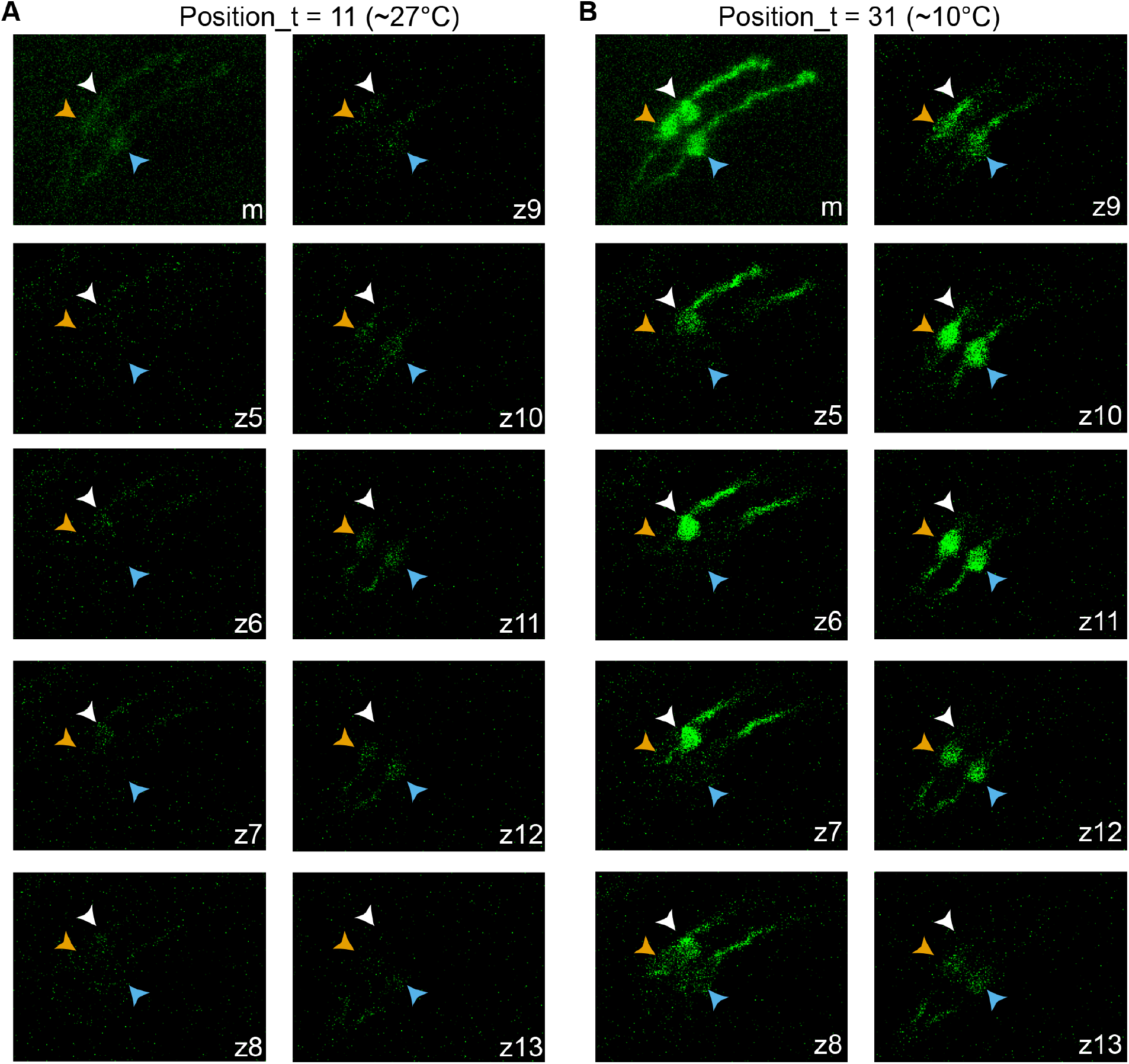
Calcium imaging of fly larval cool cells in inactive and active states. (A) Cells are barely visible in the inactive state. (B) Cells are strongly fluorescent in the active state. Different color arrowheads indicate different cells. The genotype is *Ir21a-Gal4;UAS-GCaMP6m*. m: maximal projection. z5-13: images at z-positions from 5 to 13. In (B), the cell dictated by white arrowheads is shown on z5 to z8; the cells dictated by orange and blue arrowheads are shown on z8 to z13.

**Fig 4.**
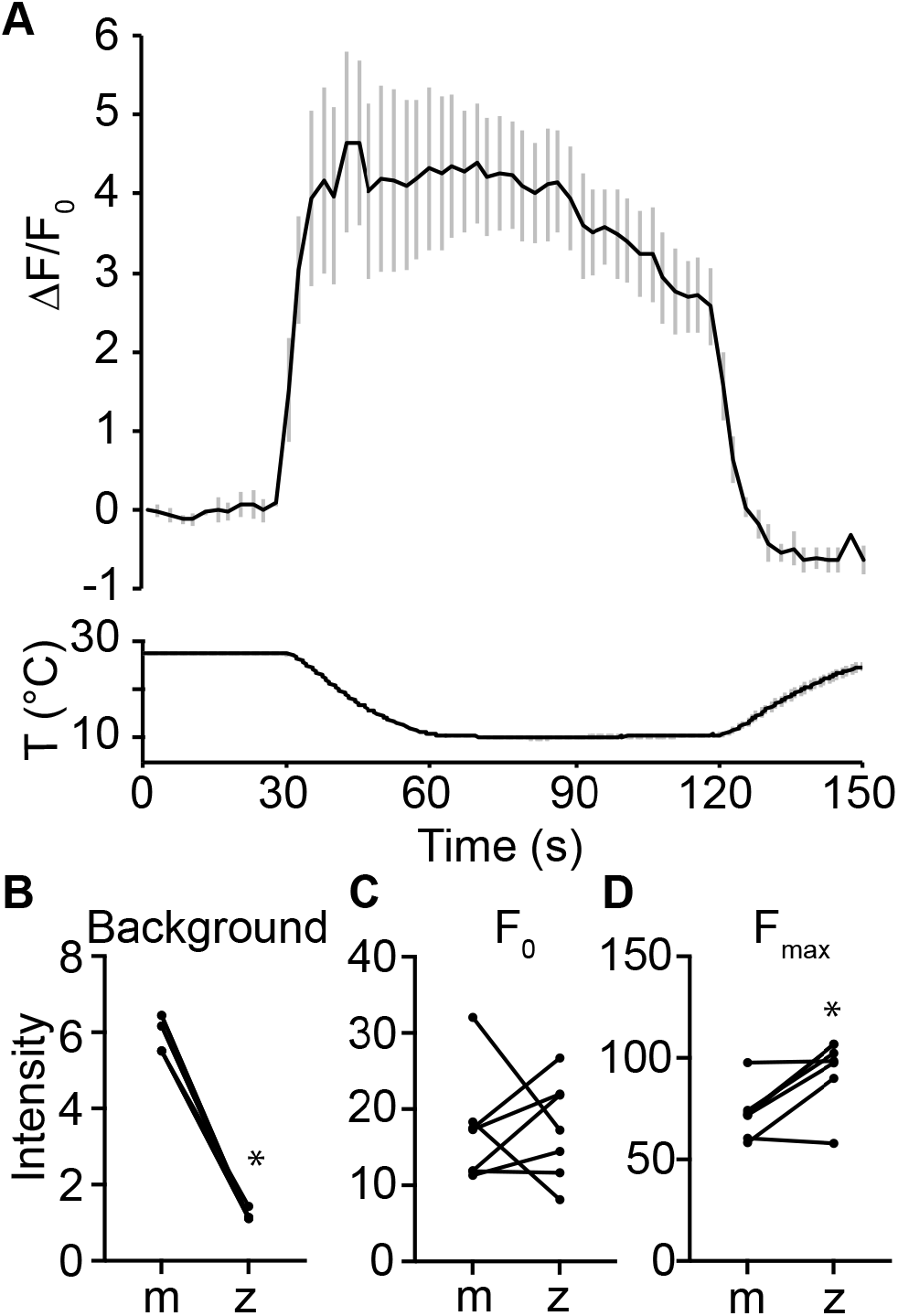
Maximal projection images overestimate background signals. (A) Fluorescence is quantified as the change in fluorescence intensity compared to the initial intensity. The genotype is *Ir21a-Gal4;UAS-GCaMP6m*. n = 7 cells from 3 animals. Traces, mean ± SEM. (B) Background intensities from maximal projection images (m) and from individual z-positions (z). Wilcoxon test, * *p* < 0.05. (C,D) Fluorescence intensities from maximal projection images (m) and from individual z-positions (z) at time points of F_0_ and F_max_. Wilcoxon test, * *p* < 0.05.

EZcalcium was also applied to analyze calcium changes to temperature fluctuations in fly larval cool neurons [12]. Although designed to correct both rigid and non-rigid motion, EZcalcium motion correction function did not always work (Fig S3A,B). Moreover, EZcalcium might extract inaccurate fluorescence intensities (Fig S3C-F).

### Maximal projection does not precisely depict the responses

In the previous study [29], maximal projection images were used to extract the fluorescence intensities (see S4_Figure in [29]). Although the trend of ΔF/F_0_ over time was similar, the plateau from maximal projection images was higher. To understand the cause of this difference, we compared the background intensities. Background intensities from individual z-positions were significantly lower than those from maximal projection images (Fig 4B). The cell intensities of F_0_ and F_max_ were also compared, and differences were random in strength and direction (Fig 4C, D). These data suggest that the extraction of fluorescence signals from maximal projection images overestimates the background intensities and may randomly affect cell signals.

### TACI separates overlapping cells

In addition, maximal projection images lose information for overlapping neurons. Fig 5A was a maximal projection image of three neurons. The white arrowhead pointed to two neurons that overlapped in the x/y plane but were separate in the ortho view (blue and orange arrowheads in Fig 5B), indicating these neurons appeared in different z-positions: the orange cell had the strongest signal on z7 (Fig 5C), while the blue cell had the strongest signal on z10 (Fig 5D). TACI distinguished these two cells and revealed the delayed but strong activation of the orange cell (Fig 5F). This information was overlooked when maximal projection images were used (Fig 5E).

**Fig 5.**
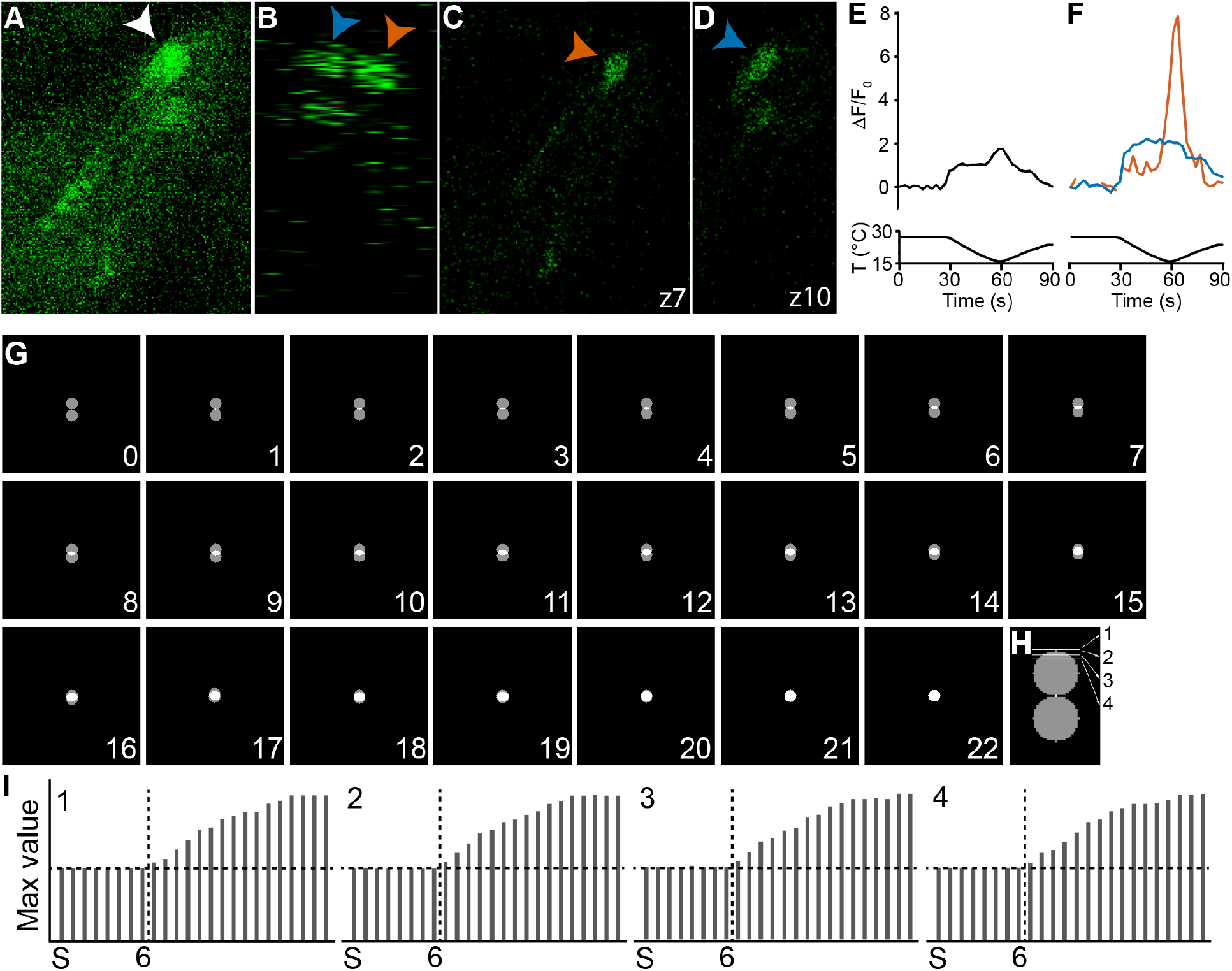
Maximum value performs better in separating overlapping cells. (A) Two neurons are overlapped (white arrowhead) in a maximal projection. The genotype is *Ir21a-Gal4;UAS-GCaMP6m*. (B) These neurons are separate in the ortho view (blue and orange arrowheads). (C,D) The orange cell appears on z7 (C), while the blue cell appears on z10 (D). (E) Fluorescence change is quantified using maximal projection images. (F) Fluorescence changes of orange and blue cells are quantified using maximum values from individual z-positions. (G) Two simulated overlapping cells in the z-axis. Each cell has a radius of 11 pixels, and each step moves one pixel, so there are 22 steps from no overlap to complete overlap. (H) Four different z-positions with a z-distance of four pixels are simulated. (I) Each z-position is analyzed. S: the maximum intensity value from a single cell. 6: overlap six pixels in (G). Horizontal dash line: the single cell’s maximum value. Vertical dash line: maximum values of the two cells are higher than that of a single cell. If the maximum value of two overlapping cells matches that of a single sphere, the distance is sufficient to identify singular spheres. Otherwise, the distance is insufficient to separate two cells.

Since the blue and orange neurons were separated in z-axis, it was possible to generate maximal projection images for each of them (Fig S4). However, it was arbitrary to determine the neuron from which the signals on the intermediate z-positions came (yellow arrowhead in Fig S4E). Importantly, when two neurons overlapped in the z-axis, it became impossible to precisely generate maximal projection images for each neuron.

To investigate whether TACI could distinguish overlapping neurons in the z-axis, we created two cells with a radius of 11 pixels. 22 one-pixel steps were required for these two cells to move from no overlap to complete overlap (Fig 5G). Four z-stacks with varied z-positions were simulated with a z-distance of four pixels (Fig 5H). If the maximum value of two cells matched the maximum value of a single cell, the distance was regarded as adequate to identify singular spheres. Otherwise, the distance was insufficient to distinguish the two cells. Our data suggest that TACI is capable of separating two cells when the overlap in the z-axis is less than half of a cell’s radius (Fig 5I).

### Analyzing a 3D calcium imaging fly brain dataset

To test whether this workflow can be used to analyze 3D calcium imaging with a large number of cells, we used a dataset acquired with a confocal microscope [11]. The imaged transgenic flies (*VT50339-Gal4;UAS-GCaMP6f*) expressed the genetically encoded calcium indicator GCaMP6f in the mushroom body in the brain [11]. Data from 45 z-positions (spaced at 1.5 μm intervals) was collected at 50 Hz for 225s (250 time points, please refer to [11] for details about preparation, equipment and experiment). We analyzed the first half of the dataset (125 time points).

When the recording began, seven neurons had obvious fluorescence, and four of them were analyzed (Fig 6A). The intensities in these neurons decreased over time (Fig 6B). When octanol was applied (Fig 6C), multiple neurons brightened. We analyzed ten neurons and found their fluorescence increased simultaneously at time point 92 (Fig 6D), suggesting that these mushroom neurons respond to octanol odor. Although octanol was applied for 5 seconds, high fluorescence in these neurons was observed in only one time point (0.9 seconds) and then quickly dropped, suggesting the response is phasic and transient. We also observed that the maximal z-drift in this dataset was 4.5 μm and the mean was 1.92 ±0.46 μm.

**Fig 6.**
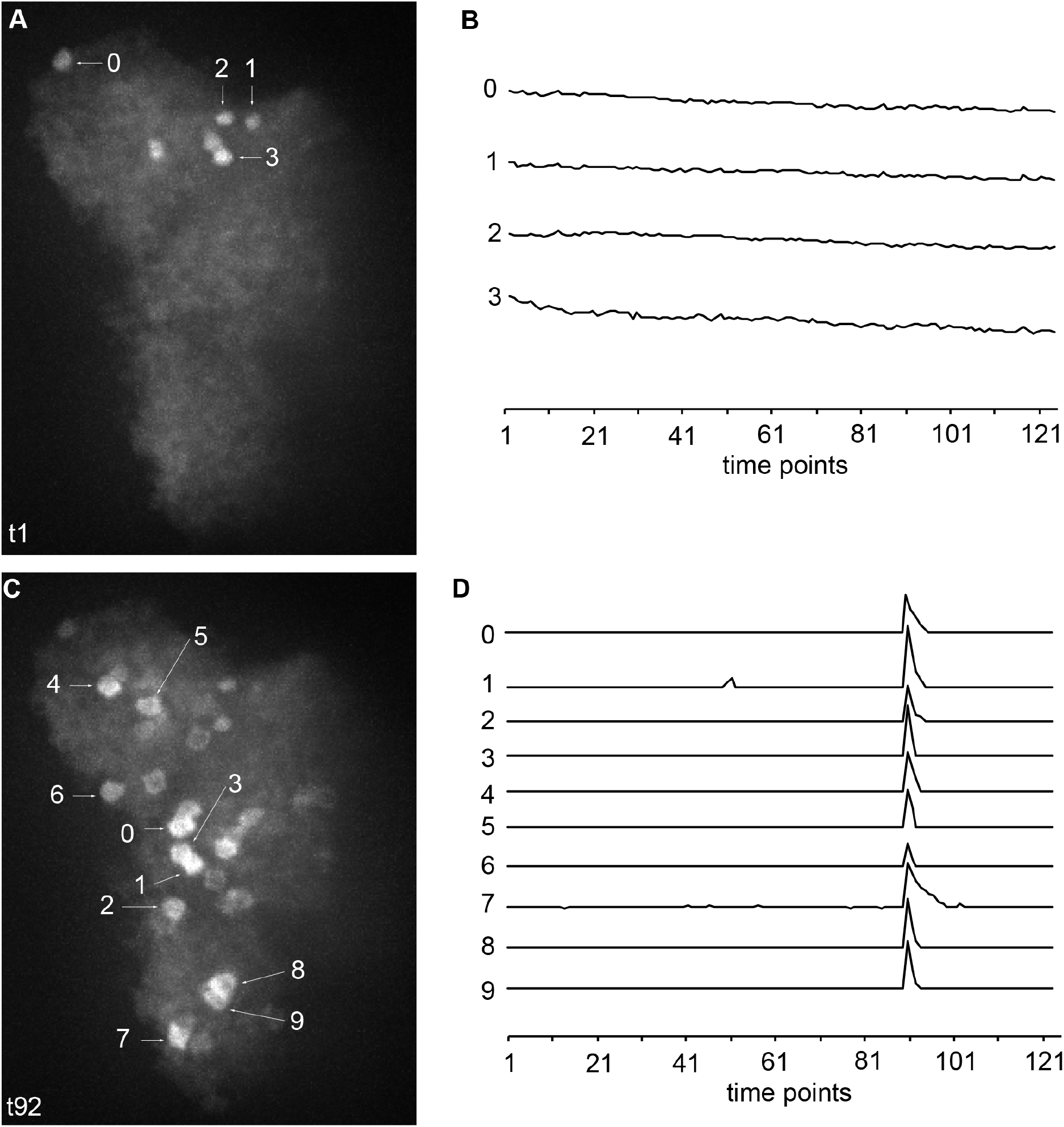
Analyzing a 3D calcium imaging fly brain dataset. (A) The maximal projection image at time point 1 (t1). 0-3 dictate the analyzed four neurons. (B) Fluorescence changes of neurons 0-3 in (A) during time points 1 to 125. (C) The maximal projection image at time point 92 (t92). 0-9 dictate the analyzed ten neurons. (D) Fluorescence changes of neurons 0-9 in (C) during time points 1 to 125.

## Discussion

This study developed a new ImageJ plugin TACI and described a workflow analyzing 3D calcium imaging and generating reproducible information of the calcium responses. Many currently available tools focus on calcium imaging data that have a large number of neurons but ignore motion on the z-axis and do not consider if individual neurons appear on multiple z-planes [6]. During image acquisition in a live organism, movement on the z-axis is unavoidable even when it is immobilized. Some stimuli, such as temperature change, often cause significant z-drift. For example, in our experiment, although animals were immobilized by cover slips, larval cool neurons still displayed a maximal z-drift of 8.25 μm and a mean of 5.25 ±0.71 μm. Increasing the height of z-stacks will record cells of interest during the whole imaging process; but it is not trivial to analyze motion on the z-axis, especially when individual cells appear on multiple z-positions. If such movement is ignored, researchers will not obtain the precise calcium responses of these cells. TACI corrects z-drift by extracting fluorescence signals from every z-position and using the maximum value to represent a cell’s intensity at each time point. It also allows for the separation of cells that partially overlap on the z-axis, and/or overlap in the lateral (x/y) direction but appear on different z-positions.

In this workflow, we used TrackMate to track cells, identify ROIs, and extract the fluorescence intensities of cells. Many studies, including a recent study from our lab [29], manually accomplished this step. This manual process is time-consuming and prone to human bias; TrackMate achieves automation of motion correction, ROI identification, and data extraction. For motion correction, we rely on TrackMate to track movements in the lateral (x/y) direction. TrackMate is designed for Brownian (random-walk) motion and receives a good evaluation for spot tracking performance [30]. This tracking-based method may be more suitable for large sudden movements than frame-based motion correction (Fig S3A,B). When using TrackMate to identify an ROI, each detected ROI is assigned a quality value – the local maximal value [20]. If this value is lower than what the detector is configured with, the ROI is discarded [20]. If an ROI cannot be detected, a decrease in the threshold makes detectors more sensitive. TrackMate creates interactive windows to allow the manual validation of every ROI. When using the same parameter settings, fluorescence intensities extracted by TrackMate are consistent from one computer or operator to another (Table S1). Another challenge for calcium analysis is that cells intermittently disappear and reappear with stimulation. TrackMate can track the cells when they reappear and register them as the same TRACK_IDs. Last but not least, TrackMate can track individual ROIs and extract their intensities from large-scale calcium imaging datasets obtained by two-photon microscopy and microendoscopy (Fig S5 and S6) [10, 31]. This workflow is therefore appropriate for analyzing calcium imaging data from different types of microscopes.

Although this workflow is semi-automatic and still requires manual efforts from researchers, it provides a computational and reproducible approach for 3D calcium imaging analysis. Importantly, this workflow is based on ImageJ and does not require programming software or knowledge. Limitations and their potential solutions for this workflow are listed below.

1. This method is not suited to analyzing a large number of neurons simultaneously. TrackMate may track over 100,000 cells. However, during image acquisition in a live organism, both non-rigid and rigid motions occur, and such movement dampens the application of TrackMate for calcium imaging in dense cells. Moreover, this workflow lacks cell registration across z-positions. Researchers must check TrackMate outcomes manually to get accurate results, and thus it becomes impractical to analyze dozens of cells. Other calcium imaging analyses can be used to replace TrackMate to correct motion and extract ROIs’ intensities. For example, we applied EZcalcium to analyze the calcium changes to temperature fluctuations in fly larval cool neurons (Fig S3).
2. Two steps in this workflow introduce unavoidable variations: (1) The background intensities may cause variation. Operators are unlikely to pick the same regions to extract background intensities. Thus, we recommend extracting background intensities from nearby same-size blobs that do not contain fluorescence signals to minimize the variation. Another recommendation is to use the average of three to five background intensities from different time points as the background intensity for the corresponding z-position. (2) Weak signals may also introduce variation. When the signals are weak, TrackMate may mistake noise as the signals of the cells. In this case, researchers must manually check and decide whether these signals are correct.
3. TACI only accepts .tif files and a specific file name structure ({file_name}_{phase}t{t}z{z}{channel}). Other file formats compatible with ImageJ can be easily converted to .tif files by ImageJ. TACI has a RENAME function that converts image names to the required structure so that TACI is compatible with calcium imaging data obtained from different systems. Calcium imaging data could be analyzed by TrackMate directly if file formats are compatible with TrackMate and the data are organized by z-positions.
4. TACI is designed for calcium imaging data with constant backgrounds. Subtracting the corresponding background information from ROI intensities at each time point is one way to correct fluctuating backgrounds. TACI provides ROI intensities at each time point in the python_files folder. The background intensities could be represented by (1) the images’ mean intensities or (2) the mean intensities of ROIs with no active cells. ImageJ provides methods to obtain the images’ mean intensities (Image>Stack>Measure Stack) (Fig S6A) and mean intensities of random ROIs (Analyze>Tools>ROI Manager>Multi Measure) (Fig S6B,C). Subtraction of mean intensities of entire images or a single spot with no active cells resulted in similar calcium changes over time (Fig S6D-O). If photobleaching happens during calcium imaging, the Bleach Correction function (Image > Adjust > Bleach Correction) may be run prior to TrackMate.
5. TACI uses the first value of each z-position as F_0_ to calculate ΔF/F_0_. If this F_0_ is not appropriate [32], TACI provides files including raw data for each neuron in the python_files folder.

## Materials and methods

### Fly strains

*Ir21a-Gal4* [28], *Ir93a-Gal4* [33]*, Ir68a-Gal4* [34], and *UAS-GCaMP6m (P{20XUAS-IVS-GCaMP6m}attp2*) [27] were previously described. *Ir21a-Gal80* was created by subcloning the *Ir21a* promoter region into *pBPGAL80Uw-6* (Addgene plasmid # 26236) [28, 35].

### Calcium imaging

Calcium imaging data in Fig 3, Fig 4, and Fig S3 were from the S4_Figure in a previous study [29] and were reanalyzed using the current workflow. The S4_Figure in the previous study [29] was analyzed using maximal projection images. Briefly, in *Ir21a-Gal4* fly larvae, dorsal organ cool neurons expressed the calcium indicator, GCaMP6 [27, 28]. Three-day-old larvae were immobilized between a glass slide and a glass coverslip with 1 x phosphate buffered saline (PBS). The temperature was held at approximately 27°C for 30 seconds and then was decreased to approximately 10°C for 30 seconds. Then, the temperature was held at 10°C for approximately 60 seconds and returned to 27°C for 30 seconds. A different temperature stimulus was applied to generate calcium imaging data in Fig 5, Fig S4, and Fig S7. In this experiment, the temperature was held at approximately 27°C for 30 seconds, decreased to approximately 15°C for 30 seconds, and then returned to approximately 24°C for 30 seconds.

### Analyzing calcium imaging

Calcium imaging files were exported to .tif files using data acquisition software or converted to .tif files by ImageJ if file formats were compatible with ImageJ. These files were then organized according to z-positions using RENAME and ORGANISE functions in TACI.

ImageJ opened all images in a folder of a single z-position and presented them as one stack in order. TrackMate was then applied to extract the fluorescence intensities of cells of interest. We recommended adjusting the following parameters in TrackMate. (1) Use DoG or LoG detectors (Fig S1A). DoG director is more sensitive than LoG when the same threshold was applied (Fig S7). (2) Change the blob diameter, threshold, and median filter (Fig S1B). Adjust the blob diameter based on the sizes of the cells. The blob diameter should be similar to the diameter of the cells. If cells were oval, the blob diameter should be similar to the minor axis. An increase in the threshold and use of the median filter helped to avoid background noise being picked up as signals. (3) Set the filters to remove some, if not all, irrelevant signals (Fig S1C). Filters X and Y were used to remove the irrelevant signals that were distant from the real signals. When filters were set on one image, it was crucial to check all other images to ensure that the real signals were not removed. We recommended analyzing them one by one. In Fig S1C, to analyze the left cell, the right cell (arrowhead) and irrelevant signal (arrow) (Fig S1C1) could be removed by setting filters X and Y (Fig S1C2). (4) Set linking max distance, gap-closing max distance, and gap-closing max frame gap (Fig S1D). We recommended setting the linking max distance and gap-closing max distance to be three to five times the blob diameter, especially when samples moved significantly over time. This setting helped decrease the number of tracks. We recommended setting gap-closing max frame gap to the number of images in the stack. (5) Export ROIs’ mean intensities (Fig S1E). If an old TrackMate version was used, choose Export all spots statistics in Select an action window (Fig S1E). If the TrackMate version was 7.6.1 or higher, choose Spots in Display options window. Both files were interactive with the image window: highlighting an ROI displayed the corresponding ROI in the image window. The same TRACK_ID was supposed to represent the same ROI at different time points. However, this was not always true and needed to be corrected manually, when necessary. These files included mean intensities (MEAN_INTENSITY or MEAN_INTENSITY_CH1) of the cell of interest at corresponding time points (POSITION_T). If TrackMate did not recognize the ROI at some time points, the time points would not be displayed.

Next, TACI EXTRACT function created a list including every time point and sorted the mean intensities into the corresponding time points, identified the maximum value of each z-stack, subtracted the background, and calculated ΔF/F_0_. TACI was designed for calcium imaging data with constant backgrounds. For calcium imaging data with fluctuating backgrounds, please refer to the Discussion section. In this study, the background intensity for each z-position was estimated by using the average value of three to five nearby same-size blobs that did not contain fluorescence signals and were from different time points. ΔF/F_0_ was calculated by the following formula. The first value of each z-position was used as F_0_.

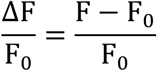

### Simulation

3D spheres with different brightness were created by a python script, and the ground truth intensities of each sphere were calculated by the following formula.

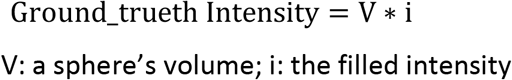

z-stacks with varied distances or positions were simulated, and TrackMate was applied to analyze these z-stacks. Ratios of maximum values or sum values and the ground truth intensities were calculated to justify that the maximum value is a good representative of a cell’s intensity.

For Fig 5, two 3D spheres of radius 11 were simulated in 3D space to depict 0 pixels overlapping to fully overlapping neurons by moving one sphere one pixel at a time. Four z-stacks with a z-distance of four pixels at different z-positions were created and analyzed by TrackMate. When the maximum value of the two spheres’ z-stack matched the maximum value of a single sphere’s z-stack, the distance was adequate to identify singular spheres. If the maximum value from the two spheres’ z-stack was higher than that from a single sphere’s z-stack, the distance was insufficient to distinguish these two spheres.

### Statistical analysis

Statistical details of experiments were mentioned in the Fig 3 legend. The normality of distributions was assessed by the Shapiro-Wilk W test (p ≤ 0.05 rejected normal distribution). For data that did not conform to a normal distribution, statistical comparisons were performed by the Wilcoxon test. Data analysis was performed using GraphPad Prism 9.

## Data and code availability

ImageJ plugin is available at: https://github.com/niflylab/TACI_CalciumImagingPlugin.

Original statistics and raw data are available at: https://doi.org/10.7910/DVN/AXEVQT.

## Acknowledgements

A Zeiss LSM 880 in the Fralin Imaging Center was used to collect calcium imaging data. We acknowledge Dr. Lenwood S.Heath for constructive comments on the manuscript and Steven Giavasis for comments on the GitHub README file. This work was supported by NIH R21MH122987 (https://www.nimh.nih.gov/index.shtml) and NIH R01GM140130 (https://www.nigms.nih.gov/) to L.N. The funders had no role in the study design, data collection and analysis, decision to publish, or preparation of the manuscript.

---

**Fig S1.**
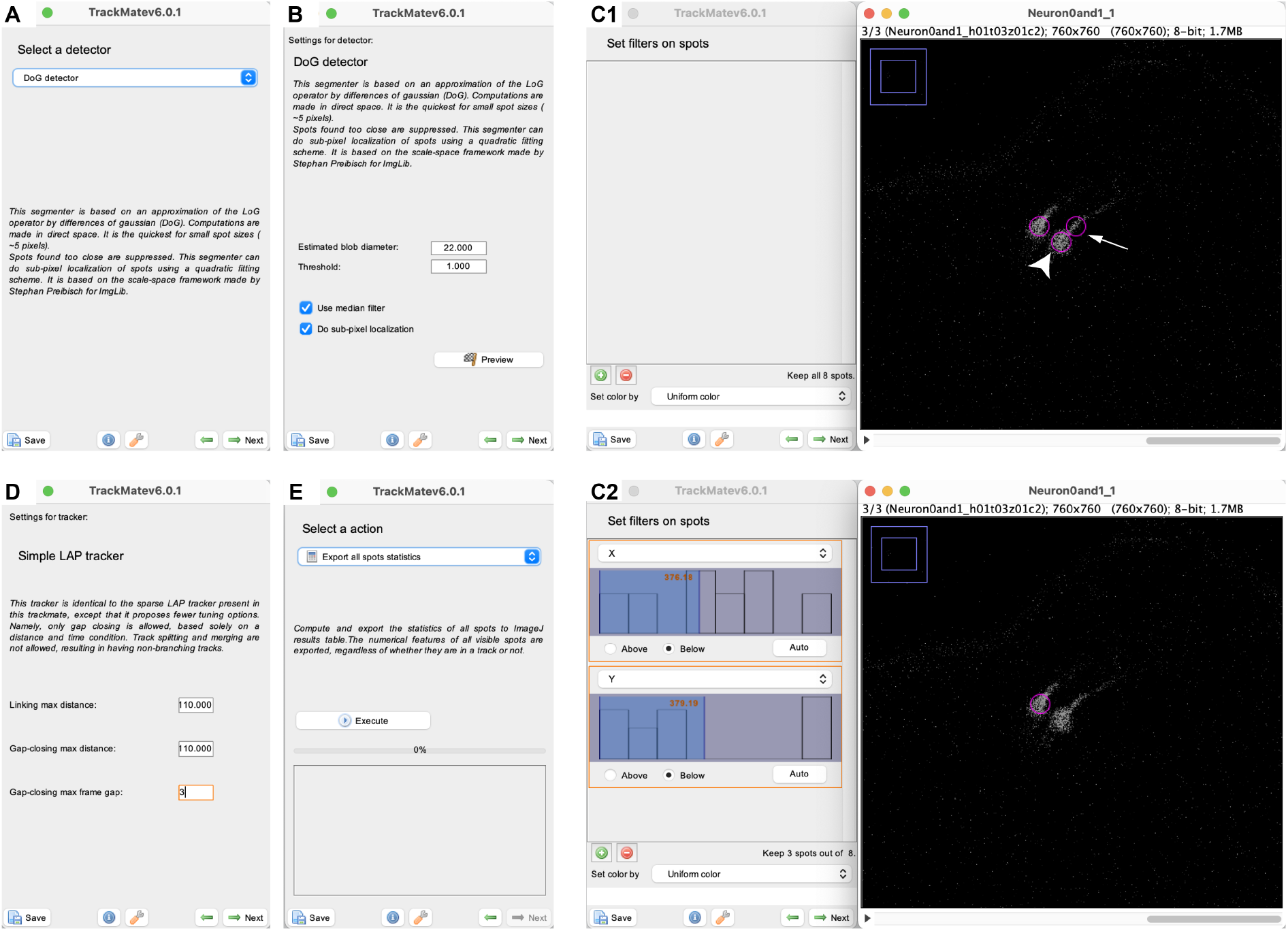
Use TrackMate to extract cells’ intensities. The following parameters are recommended to be adjusted. (A) Select a detector. (B) Set parameters of the detector. If a DoG detector is used, we recommend changing the blob diameter based on the sizes of the cells. The blob diameter should be similar to the diameter of the cells. If cells are oval, the blob diameter should be similar to the minor axis. Increasing the threshold can help decrease the effects of the background noise. If signals are strong, we recommend using the median filter, which can help decrease the Salt and Pepper noise. (C) Place filters on spots. When irrelevant signals are picked up, filters help remove some, if not all, irrelevant signals. Filters X and Y can easily remove the irrelevant signals that are distant from the real signals. When filters are placed on one image, it is crucial to check all other images to make sure the real signals are not removed. In (C1), two cells of interest and one irrelevant signal (arrow) are picked up (magenta circles). To analyze the left cell, the right cell (arrowhead) and irrelevant signal (arrow) can be removed by setting filters X and Y (C2). (D) Simple LAP tracker. We recommend setting linking max distance and gap-closing max distance to be three to five times the blob diameter, especially when samples move over time. These settings help decrease the number of tracks. We recommend setting the gap-closing max frame gap to the number of images in the folder. (E) Select an action. We recommend choosing the option of Export all spots statistics. If a new version of TrackMate is used, similar information is exported by choosing Spots in the Display options window.

**Fig S2.**
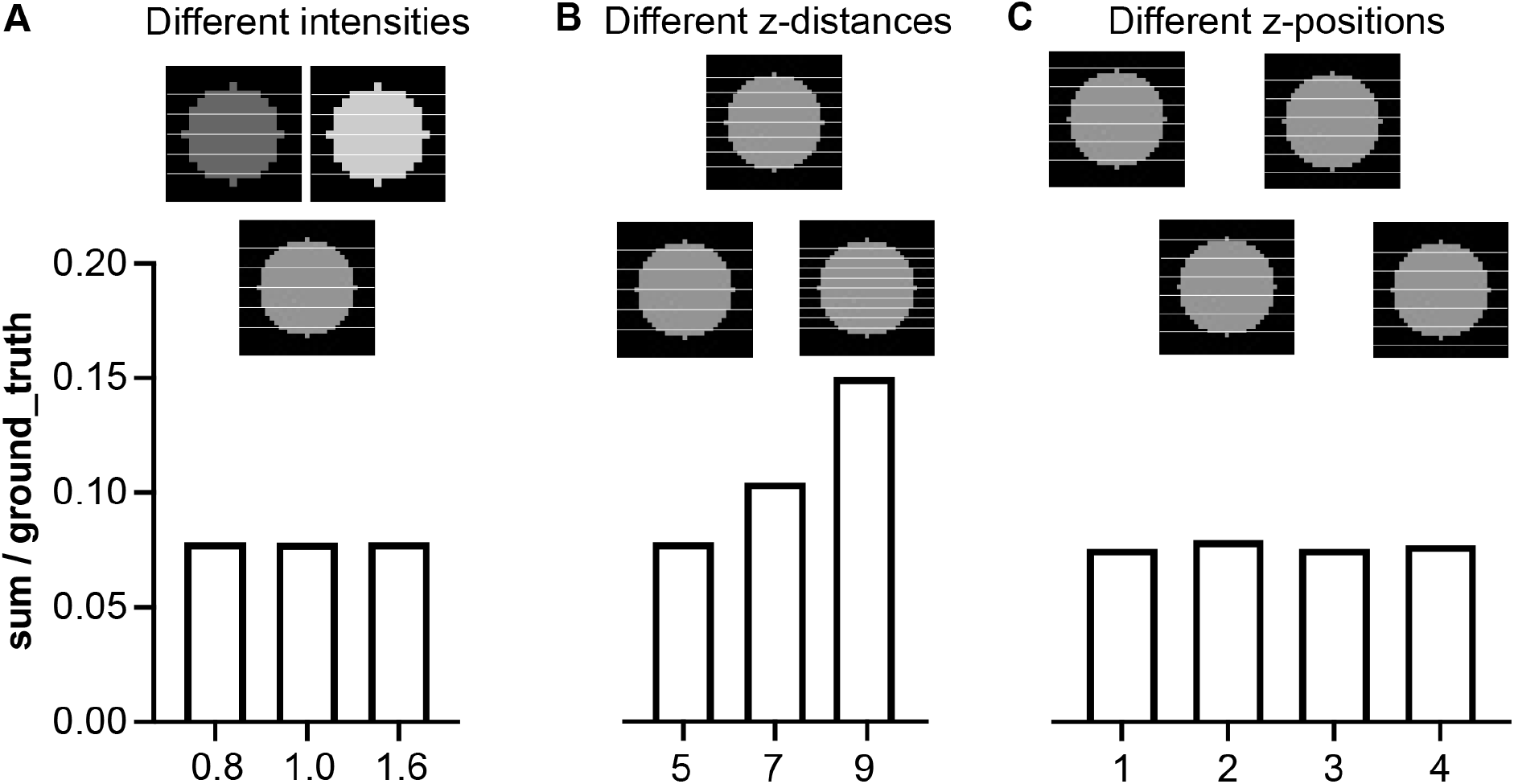
The sum value may not be a good representative of a cell’s intensity. (A) z-stacks are created from simulated cells with different intensities. (B) z-stacks with different z-distances are simulated. (C) z-stacks with different z-positions are simulated. sum: the total intensities of all z-positions. ground_truth: the product of the simulated cell’s volume and its filled intensity.

**Fig S3.**
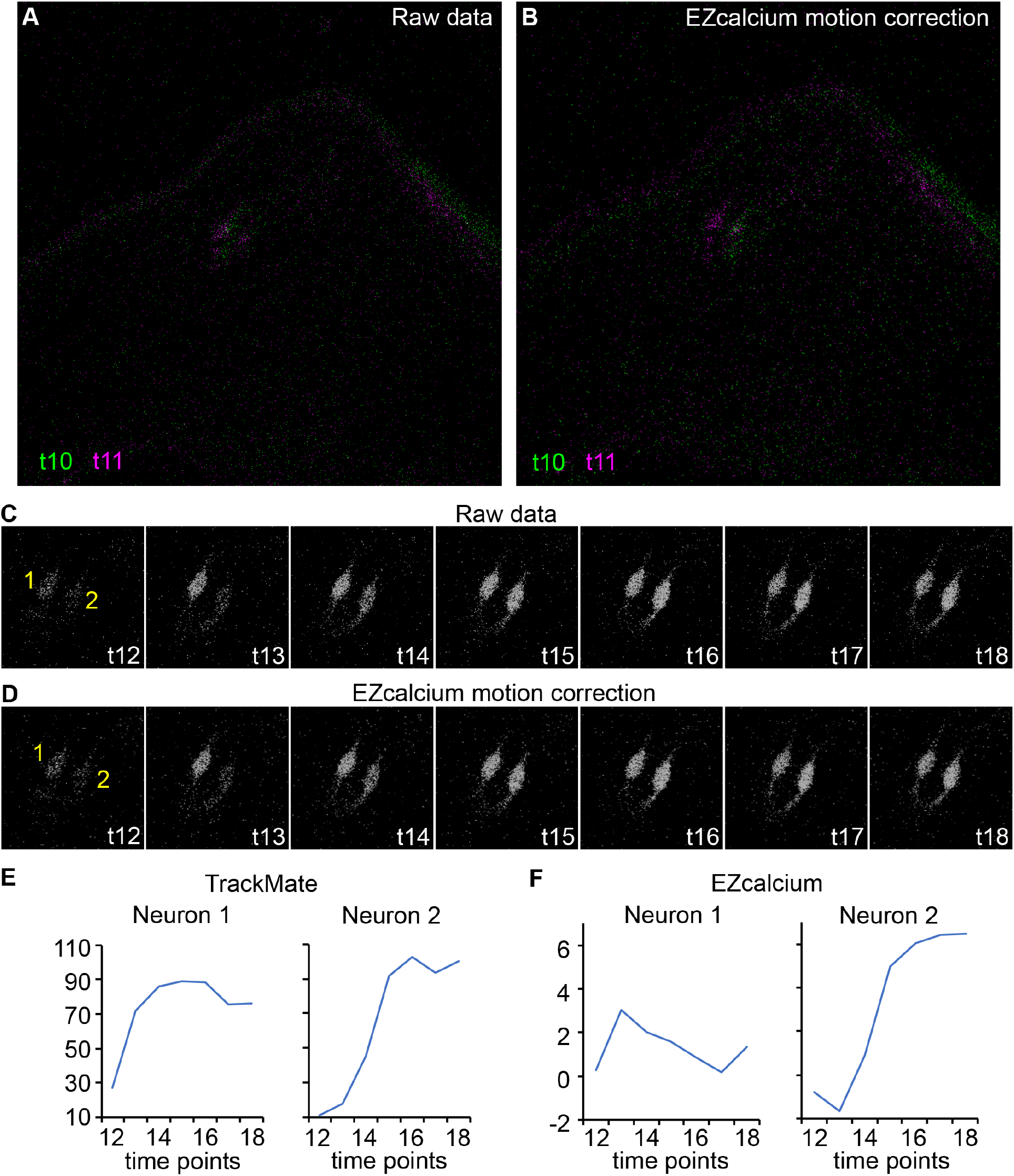
TrackMate outperforms EZcalcium in cool cell calcium imaging analysis. (A,B) EZcalcium does not always correct the x/y motion of cool cells. All possible non-rigid settings with their maximum allowed movements have been tested. The genotype is *Ir21a-Gal4;UAS-GCaMP6m*. (A) Raw images. (B) Images after motion correction by EZcalcium. Green: time point 10 (t10). Magenta: time point 11 (t11). Images are at z-position 8. (C,D) Seven continuous time points (t12-18) from raw images (C) and images after motion correction by EZcalcium (D). Yellow 1 and 2 indicate Neuron 1 and Neuron 2, respectively. (E) Fluorescence intensities are extracted by TrackMate from (C). (F) Fluorescence intensities are extracted by EZcalcium from (D). Images are at z-position 8. Of note, EZcalcium automatedly recognizes neither neuron and thus manual ROI selection is performed. The ROI shapes are determined by EZcalcium.

**Fig S4.**
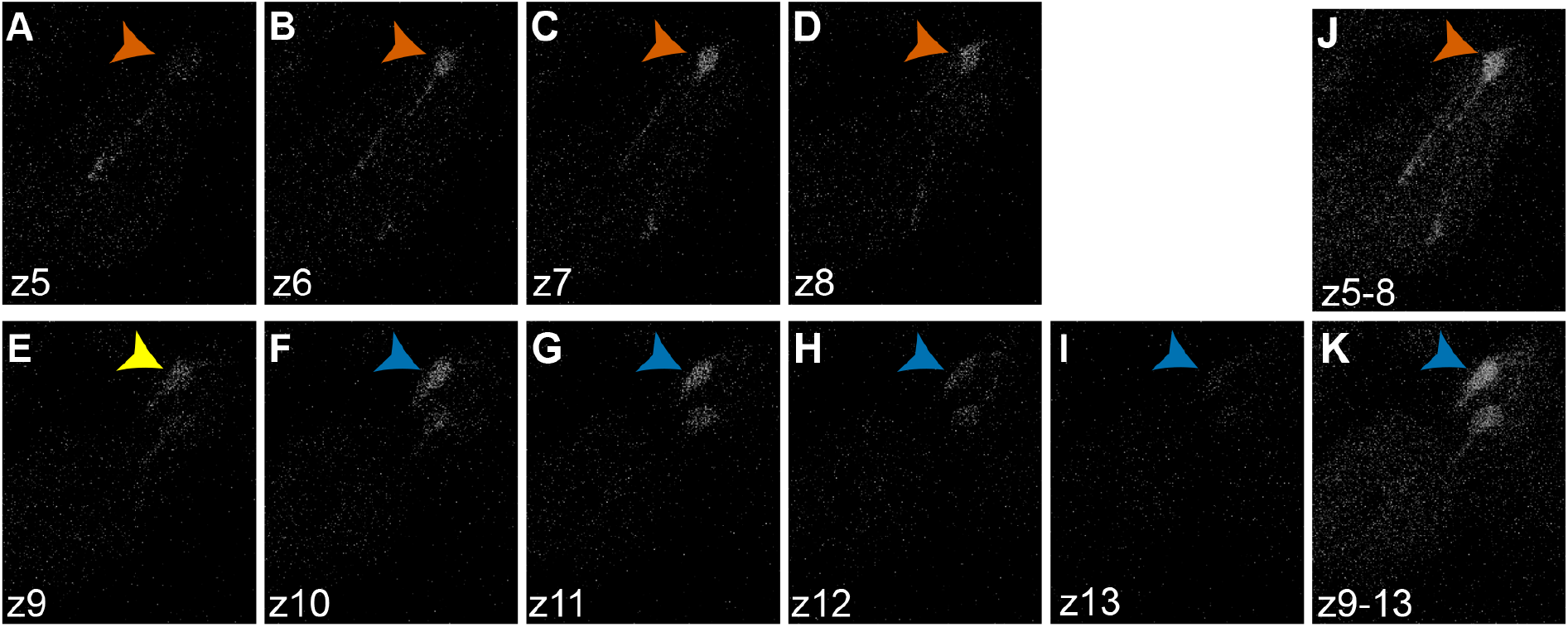
Generation of maximal projection images of overlapping cells. (A-I) z-positions from 5 to 13. (J) The maximal projection image from z5 to z8. (K) The maximal projection image from z9 to z13. The same calcium imaging data as **Fig 5**. Of note, it is subjective to decide whether the signal at z9 (yellow neuron in (E)) is from the orange cell or the blue cell.

**Fig S5.**
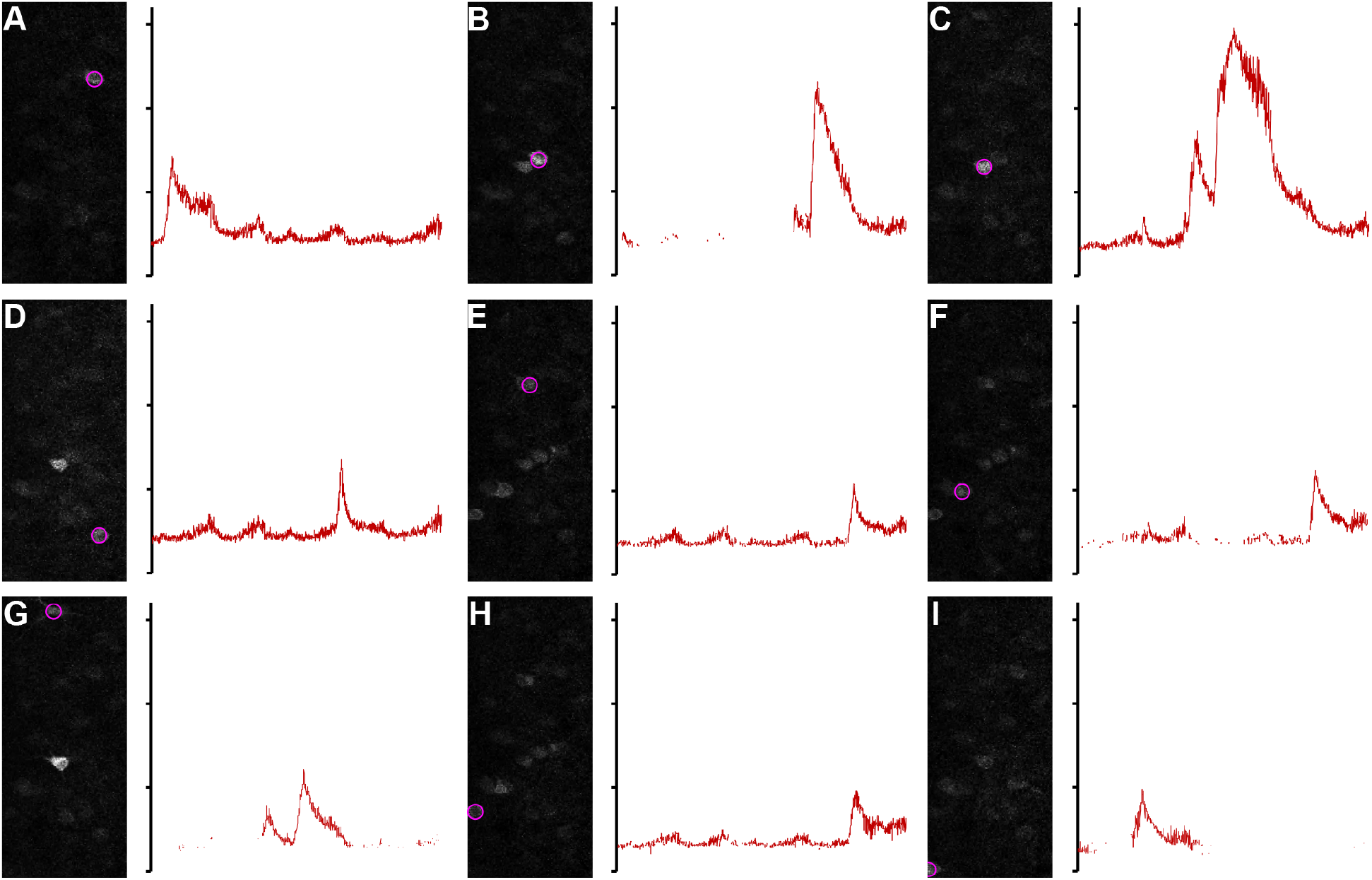
TrackMate analyzes two-photon calcium imaging of hippocampal activity. Fluorescence changes of nine neurons (magenta in A-I) are analyzed by TrackMate. The y-axis of the right panel is fluorescence intensities extracted by TrackMate of a range between 0 and 160.

**Fig S6.**
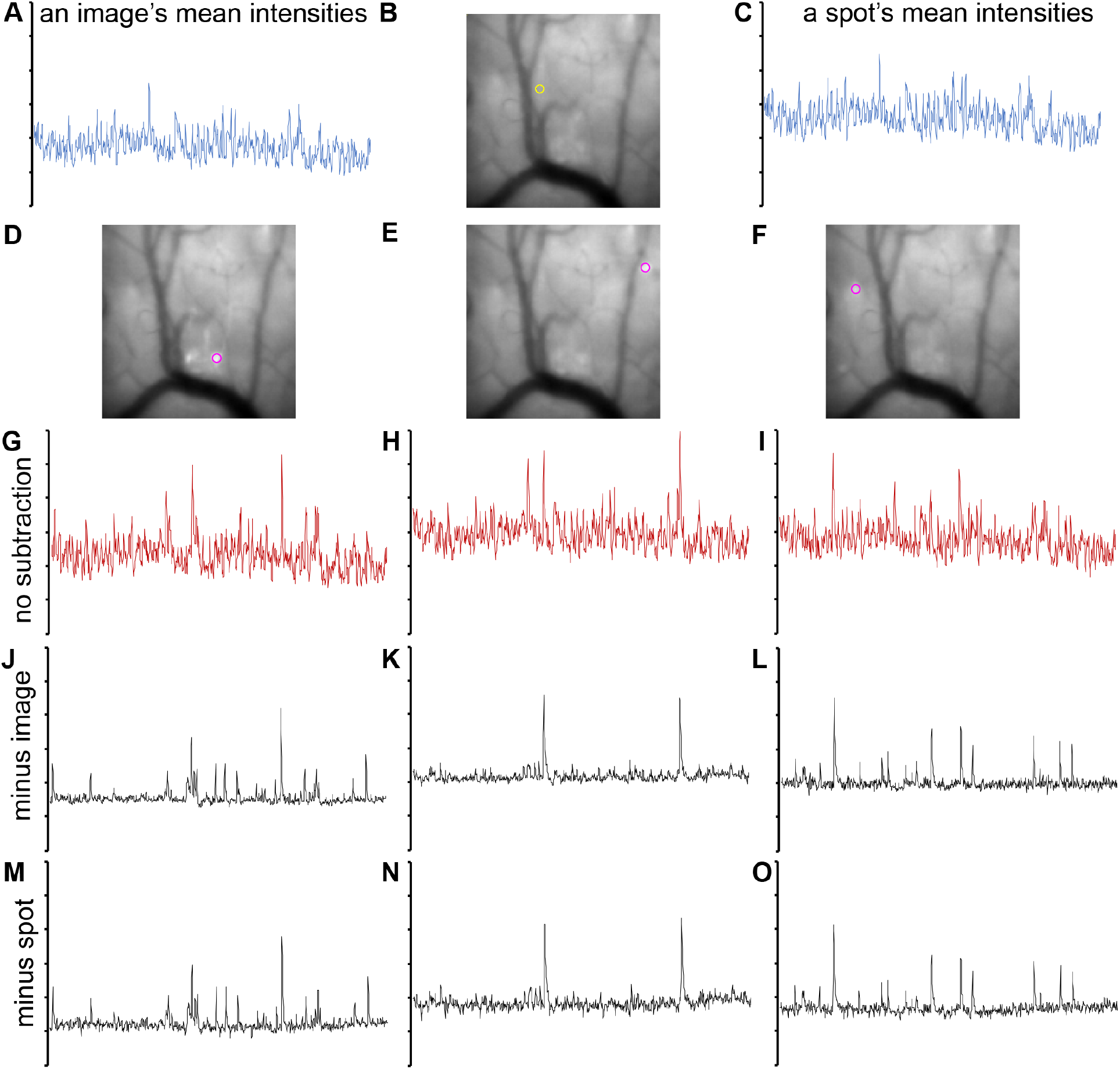
TrackMate analyzes microendoscopic data recorded in the dorsal striatum. (A) The mean intensities of each image are extracted by TrackMate. y-axis: fluorescence intensities (90-210). (B,C) Mean intensities of a spot (yellow in B) are extracted from every image by TrackMate. This spot does not include active neurons during the whole recording process. y-axis: fluorescence intensities (110-230). (D-O) Fluorescence changes of three neurons (magenta in D-F) are analyzed by TrackMate. (G-I) Mean intensities of the corresponding neurons in D-F are extracted by TrackMate. y-axis: fluorescence intensities (110-230). (J-L) Fluorescence changes are indicated by the difference between neurons’ intensities (G-I) and mean intensities of whole images (A). y-axis: fluorescence intensities (0-120). (M-O) Fluorescence changes are indicated by the difference between neurons’ intensities (G-I) and a non-active spot’s intensities (C). y-axis: fluorescence intensities (−30-90).

**Fig S7.**
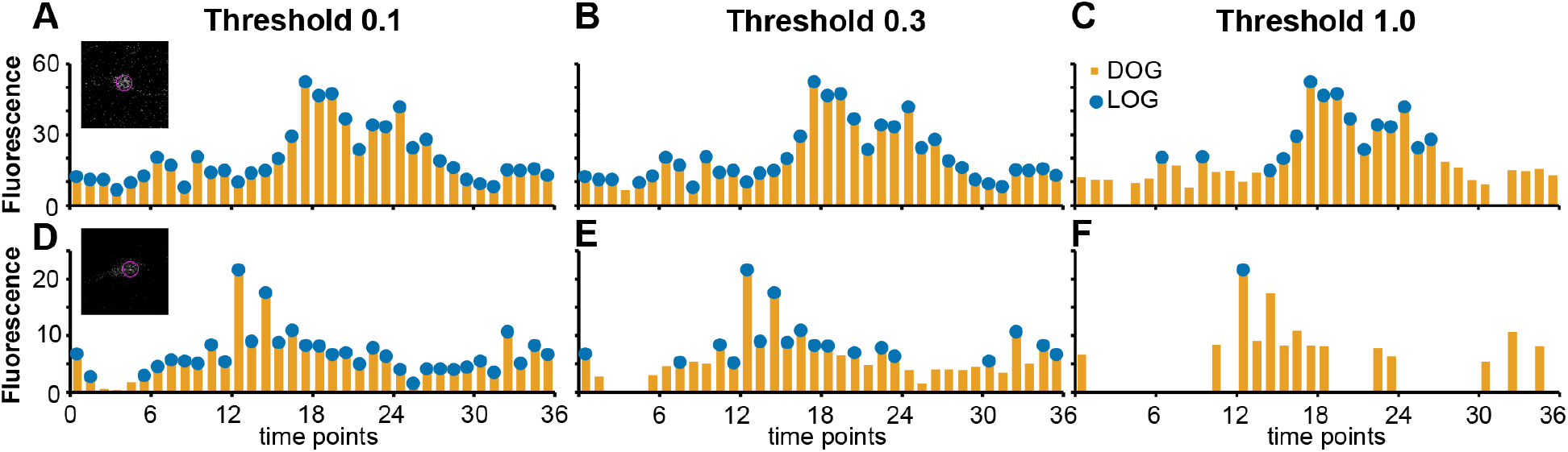
The DoG detector is more sensitive to weak signals. Two neurons with weak calcium signals are analyzed. (A-C) The first neuron (*Ir21a-Gal80/UAS-GCaMP6;Ir93a-Gal4*) is analyzed by DoG and LoG at different thresholds. (A) The threshold is set at 0.1. Both DoG and LoG detect all 36 time points. (B) The threshold is set at 0.3. Among 36 time points, DoG detects 36, and LoG detects 35. (C) The threshold is set at 1.0. Among 36 time points, DoG detects 34, and LoG detects 15. (D-F) The second neuron (*UAS-GCaMP6;Ir68a-Gal4*) is analyzed by DoG and LoG at different thresholds. (D) The threshold is set at 0.1. DoG detects all 36 time points, and LoG detects 34. (E) The threshold is set at 0.3. Among 36 time points, DoG detects 33, and LoG detects 18. (F) The threshold is set at 1.0. Among 36 time points, DoG detects 14, and LoG detects only one.

**Table S1. TrackMate generates reproducible results.** Two neurons are tested by three operators, four computers (three Mac and one Windows), and two TrackMate versions. When TrackMate versions and parameters are the same, different operators and computers export the same results. If using different TrackMate versions, the results are slightly varied even with the same parameters. Yellow cells indicate different results from the two TrackMate versions.

**Movie S1. A virtual tutorial to explain how to use TACI.**

